# Identification of novel components of the Ced and Ups systems in *Saccharolobus islandicus* REY15A

**DOI:** 10.1101/2024.09.27.615246

**Authors:** Pengju Wu, Mengqi Zhang, Yanlu Kou, Shikuan Liang, Jinfeng Ni, Qihong Huang, Yulong Shen

## Abstract

In the cells of *Sulfolobales*, the Ced (Crenarchaeal system for exchange of DNA) and Ups (UV-inducible pili of *Sulfolobus*) systems are highly induced by DNA damages and function in chromosomal DNA exchange by mediating DNA import and cell aggregation, respectively. The Ced system is composed of CedA, CedA1, CedA2, and CedB. The Ups is composed of UpsA, UpsB, UpsE and UpsF. However, how the DNA is transported by these systems is far from clear. Here, we report two novel components of the Ced system, CedD (SiRe_1715) and CedE (SiRe_2100), which are paralogs of CedB and CedA, respectively, and one new component of the Ups system, UpsC (SiRe_1957), which is a paralog of UpsA and UpsB, in *Saccharolobus islandicus* REY15A. DNA exchange assay revealed that *cedD*, *cedE*, and *upsC* are essential for DNA import while *cedE* and *upsC* are also important for DNA export. Further microscopy analysis revealed that *upsC* is involved in cell aggregation like other Ups genes. Besides, we found that *cedB* and *cedD* co-occur in the genomes of crenarchaea which lack *virB4* and the presence of *cedB* and *cedD*, and *virB4* is complementary. Both proteins show homology to different parts of the N-terminal domain of VirB4 in a complementary manner and form stable homo-oligomers *in vitro*. Collectively, our results indicate that CedD and CedE, and UpsC are essential components of the Ced and Ups systems, respectively.

## Introduction

Horizontal gene transfer (HGT) is essential for the adaption of life to the environments. In archaea, HGT includes classical mechanisms, natural transformation, conjugation, and transduction (Wagner *et al*., 2017), and archaea-specific mechanisms, such as DNA transferred by extracellular vesicles in *Thermococcales* and *Sulfolobales* (Gaudin *et al*., 2013; Liu *et al*., 2021), and chromosomal DNA exchange in haloarchaea and *Sulfolobales* (Mevarech & Werczberger, 1985; Grogan, 1996). In the hyperthermophilic crenarchaea of the order *Sulfolobales*, the inter-cellular chromosomal recombination frequency is 10^-4^∼10^-5^, much higher than that of spontaneous mutation (1-3×10^-7^), suggesting that the cells of *Sulfolobales* utilize efficient DNA transfer to overcome frequent DNA damages under high temperature (Jacobs & Grogan, 1997; Ghane & Grogan, 1998). Further, it is established that chromosomal DNA exchange in *Sulfolobales* is simulated by UV-irradiation and cell aggregation mediated by Ups (UV-inducible pili of *Sulfolobus*) is indispensable for the DNA exchange (Schmidt *et al*., 1999; Ajon *et al*., 2011). Subsequently, a system called Ced (Crenarchaeal system for exchange of DNA) was found to be involved in DNA import (van Wolferen *et al*., 2016). However, how the DNA is transported by the Ced and Ups systems remains unclear.

Ced system is composed of four transmembrane (TM) proteins, CedA, CedA1, CedA2, and CedB (van Wolferen *et al*., 2016). CedA and CedB are homologs of bacterial VirB6 and VirB4, respectively, and function in DNA import but not export (van Wolferen *et al*., 2016). CedA1 and CedA2 are composed of only two hydrophobic helixes and their functions in DNA transport have not been identified. Recently, it has been reported that the CedA1 homolog from *Aeropyrum pernix* K1 forms T4SS (Type Ⅳ secretion system)-like pili, suggesting that CedA1 could function as a conduit for DNA transport (Beltran *et al*., 2023). T4SS generally contains VirB2-VirB11 and VirD4 components (Li & Christie, 2018; Yuan *et al*., 2018; Mace *et al*., 2022). The main function of T4SS is to export conjugative DNA or effectors (Backert *et al*., 2000; Llosa *et al*., 2002). In the process of conjugation, double-stranded conjugative plasmid is firstly processed by a relaxase complex to form ssDNA which is then delivered to the ATP-powered components, VirD4, VirB11, and VirB4 sequentially (Atmakuri *et al*., 2004; Waksman, 2019). The DNA export machinery VirB2-VirB11 complex is composed of IMCC (Inner membrane core complex), OMCC (Outer membrane core complex), and the stalk which connects the IMCC and OMCC (Mace *et al*., 2022). The ssDNA is exported by the VirB4 ATPase ring which is composed of six VirB4 dimers (Mace *et al*., 2022).

Then, the ssDNA is delivered through the transmembrane channel formed by VirB6 and enters the F-pili formed by VirB2 (Jakubowski *et al*., 2004; Judd *et al*., 2005; Goldlust *et al*., 2023; Beltran *et al*., 2024). In rare case, the T4SS system (such as the ComB system in *Helicobacter Pylori*) is used to uptake DNA (Stingl *et al*., 2010). So far only the homologs of VirB2, VirB4, and VirB6 of the bacterial T4SS system are found in the Ced system (Beltran *et al*., 2023). This raises questions about how DNA import is fulfilled by the archaeal Ced system and if there are additional components of the Ced system.

Different from archaeal conjugation in which conjugative plasmids can even be transferred from *Thermococcales* to *Desulfurococcales* (Catchpole *et al*., 2023), the chromosomal DNA exchange in *Sulfolobales* is species-specific (Ajon *et al*., 2011). This species-specific recognition is mediated by Ups, one of the T4Ps (Type Ⅳ pili) (van Wolferen *et al*., 2020). In *Saccharolobus islandicus* REY15A, there are four kinds T4Ps, archaellum, adhesion pili, Ups, and putative bindosome (Szabo *et al*., 2007; Zolghadr *et al*., 2007; Frols *et al*., 2008; Henche *et al*., 2012). Only Ups is induced by DNA damages and mediates cell aggregation (Frols *et al*., 2008). The genes coding for Ups components are encoded by a conserved *ups* operon and are arranged as *upsX*-*upsE*-*upsF*-*upsA*-*upsB* in all *Sulfolobales* except *Acidianus*, in which the *upsF*-*upsA*-*upsB* are missing (van Wolferen *et al*., 2013; van Wolferen *et al*., 2016). UpsX is a membrane protein with unclarified function and deletion of *upsX* did not affect cell aggregation (van Wolferen *et al*., 2013). UpsE and UpsF function as the pilin assembly ATPase and platform, respectively, while UpsA and UpsB are pilin subunits forming pilus which bridges species-specific aggregation by recognition of different glycosylation patterns (van Wolferen *et al*., 2013; van Wolferen *et al*., 2020).

Interestingly, many uncharacterized genes are highly upregulated together with those encoding for the Ced and Ups systems when the cells of *Sulfolobales* are exposed to DNA damage agents (Frols *et al*., 2007; Feng *et al*., 2018; Schult *et al*., 2018; Sun *et al*., 2018). Some of these genes could be candidates encoding new components for DNA transport (import and export) as well as intercellular contact and recognition. In this study, we report the identification and bioinformatics analysis of three new components designated as CedD, CedE, and UpsC of the Ced and Ups systems. By chromosomal DNA exchange assay and cell aggregation analysis, we found that CedD is only involved in DNA import, probably by forming a DNA translocation channel with CedB, and CedE is involved in both DNA export and import, while UpsC is a new pilin subunit essential for cell aggregation.

## Materials and methods

### Strains and culture conditions

*Sa. islandicus* REY15A (E233) (Δ*pyrEF*) and its derived strains are shown in Supplementary Table 1. The cells were cultured at 75℃ in MTVU media supplemented 0.2 % (w/v) D-arabinose or sucrose as previously reported (Huang *et al*., 2020). Knockout of genes encoding for the Ced system, Ups, and other T4Ps was conducted by transformation of the knockout plasmids (Supplementary Table 2) into E233S (Δ*pyrEF*Δ*lacS*) followed by selection of the transformants as the previous study (Li *et al*., 2016).

### CRISPR-Cas based chromosomal DNA export and import assay

The chromosomal DNA exchange assay was modified from the DNA import assay described previously (Liu *et al*., 2020). Efficient targeting protospacers for *lacS* and *amyα* (Supplementary Table 3) were selected and cloned into the mini-CRISPR array of pGE (Li *et al*., 2016), yielding pT*lacS* and pT*amyα*, respectively. For DNA export assay, 1 μg pT*lacS* plasmid was electroporated into E233 (with *lacS*) cells (approximately 10^8^) and the cells were incubated in 0.5 mL preheated medium containing mineral salts (M) to recover. At the same time, 10^8^ competent cells of the donor strains, E233S (*lacS* deleted) or its derivates Δ*cedD*, Δ*cedE*, or Δ*upsC*, etc., were incubated in 0.5 mL preheated 2×MTV medium containing 0.4 % (w/v) D-arabinose at 75℃. After 1 hour, the two cultures were mixed and incubated at 75℃ for 2 h to exchange DNA. Then, a total of the 1 mL culture was plated on MTAV plate and cultured for 8 to 9 days. The DNA export efficiency was defined as the colony forming unit (CFU) divided by the transformation efficiency of E233. For the DNA import assay, 1 μg pT*amyα* plasmid was electro-transformed into the receptor strain, E233S or its derivates Δ*cedD*, Δ*cedE*, or Δ*upsC*, etc., which contains the complete *amyα*. The cells were mixed and incubated with the donor strain Δ*amyα* following the same steps as those for the DNA export assay. The DNA import efficiency was defined as the CFU divided by the transformation efficiency of each receptor strain. Finally, relative DNA export and import efficiency of the strains was normalized to the efficiency of E233S as 100 %.

### Phylogenetic analysis and structural prediction

For phylogenetic analysis, the homologs of CedB, CedD, and VirB4 were searched in the genome of the representative Crenarchaea species by BLAST. Then the sequences were aligned by MAFFT 7.0 Webserver. Sequences of the ATPase domains of a total of 88 homologs from 47 species were analyzed by MEGA 7.0 using the Maximum Likelihood method based on the LG+G+I+F model (Le & Gascuel, 2008) and annotated by iTOL (https://itol.embl.de/). The homologs of CedA, CedE, and VirB6 in archaea were searched by AlphaFold Cluster (Barrio-Hernandez *et al*., 2023) and the sequence were directly exported from these accessions (A0A2T9WR39, A0A256ZYB0, and A0A7C2LFF1). The phylogenetic tree was constructed using the same steps and based on LG+G+F model. The structural prediction was performed by ColabFold and AlphaFold server using default parameters (Mirdita *et al*., 2022). The structural homologs were searched by Foldseek Search (van Kempen *et al*., 2023) and analyzed by PyMOL.

### UV treatment and RT-qPCR analysis

The culture (30 mL) of *Sa. islandicus* E233S were treated with 100 J/m^2^ UV at OD_600_=0.2 and cultured at 75℃ in the dark. The samples were taken at 0 h, 3 h, 6 h, 12 h, and 24 h after treatment. Total RNA extraction, cDNA synthesis, and qPCR were performed as previously reported (Yang *et al*., 2023).

### Protein expression and purification

The C-terminal His-tagged CedBΔTM and C-terminal Flag-tagged CedDΔTM co-expression strain E233SΔ*cedB*Δ*CedD*/piDSB-C-His-CedBΔTM/C-Flag-CedDΔTM was cultured to OD_600_=0.3 in 1 L MTV medium. Then 10 mL 20 % (w/v) D-arabinose was added to induce protein expression and DNA double-strand breaks for 2 days. About 4 L culture were collected and resuspended in a lysis buffer (50 mM pH 8.0 Tris-HCl, 400 mM NaCl, 5 % glycerol). After sonication, the sample was centrifuged at 4℃, 13,000 *g* for 30 min and the supernatant was loaded onto a Ni-NTA column and the flowthrough was collected. Then the column was washed with the wash buffer (20 mM iminazole in the lysis buffer) and eluted with the elution buffer (400 mM iminazole in the lysis buffer). The flowthrough which contains Flag-tagged CedDΔTM was further purified using anti-Flag magnetic agarose. The eluted fractions of nickel column and anti-Flag magnetic agarose were concentrated by ultrafiltration and then loaded onto a Superdex 200 increase 10/300 (GE healthcare) column, respectively. Size exclusive chromatography (SEC) was conducted using the lysis buffer and the fractions were analyzed by SDS-PAGE, Coomassie blue staining, and Western blotting as described previously (Yang *et al*., 2023).

### Cell aggregation analysis

Strains Δ*cedA*ΔFAB, Δ*upsC*ΔFAB, Δ*upsXEFAB*ΔFAB, and the control ΔFAB, in which the genes for archaellum, adhesion pili, and bindosome were deleted, were subjected to cell aggregation analysis. After UV treatment, the samples were taken as described above and the cell morphology was examined by microscopy under a NIKON TI-E inverted fluorescence microscope (Nikon, Japan) in differential interference contrast (DIC). The aggregation was defined as the aggregates containing at least 3 cells.

## Results

### Identification of putative genes involved in DNA transfer

Previously, the Ced and Ups systems were identified by analyzing the highest upregulated genes after UV irradiation in *Sulfolobales* (Frols *et al*., 2007; Frols *et al*., 2008; van Wolferen *et al*., 2016). To find additional factors that are involved in DNA transfer, we reannotated the rest 14 genes with more than 30-folds upregulation after DNA damage agent (NQO) treatment in *Sa. islandicus* REY15A (Sun *et al*., 2018) by BLAST, Foldseek, and AFDB clusters (Supplementary Table 4). We focused on three genes, *sire_1715*, *sire_2100*, and *sire_1957* and named the encoded proteins as CedD, CedE, and UpsC, respectively, based on the results from this study. CedD is composed of a TM domain, a functionally unknown N-terminal domain (NTD), and a HerA/VirB4/CedB-like ATPase domain at the C-terminal (CTD) (Figure 1A). CedE, a CedA paralog, was only found in *Saccharolobus* (Supplementary Figure 1). It contains five transmembrane (TM) helixes and has structural similarity to CedA and bacterial VirB6 (Figure 1B). UpsC is a prepilin domain-containing protein, in which the N-terminal domain and part of the C-terminal domain can align with UpsB and UpsA, respectively (Figure 1C). UpsC is only conserved in *Sulfolobales* (except in the genus *Acidianus*) like the reported Ups system proteins (van Wolferen *et al*., 2016).

**Figure 1.**
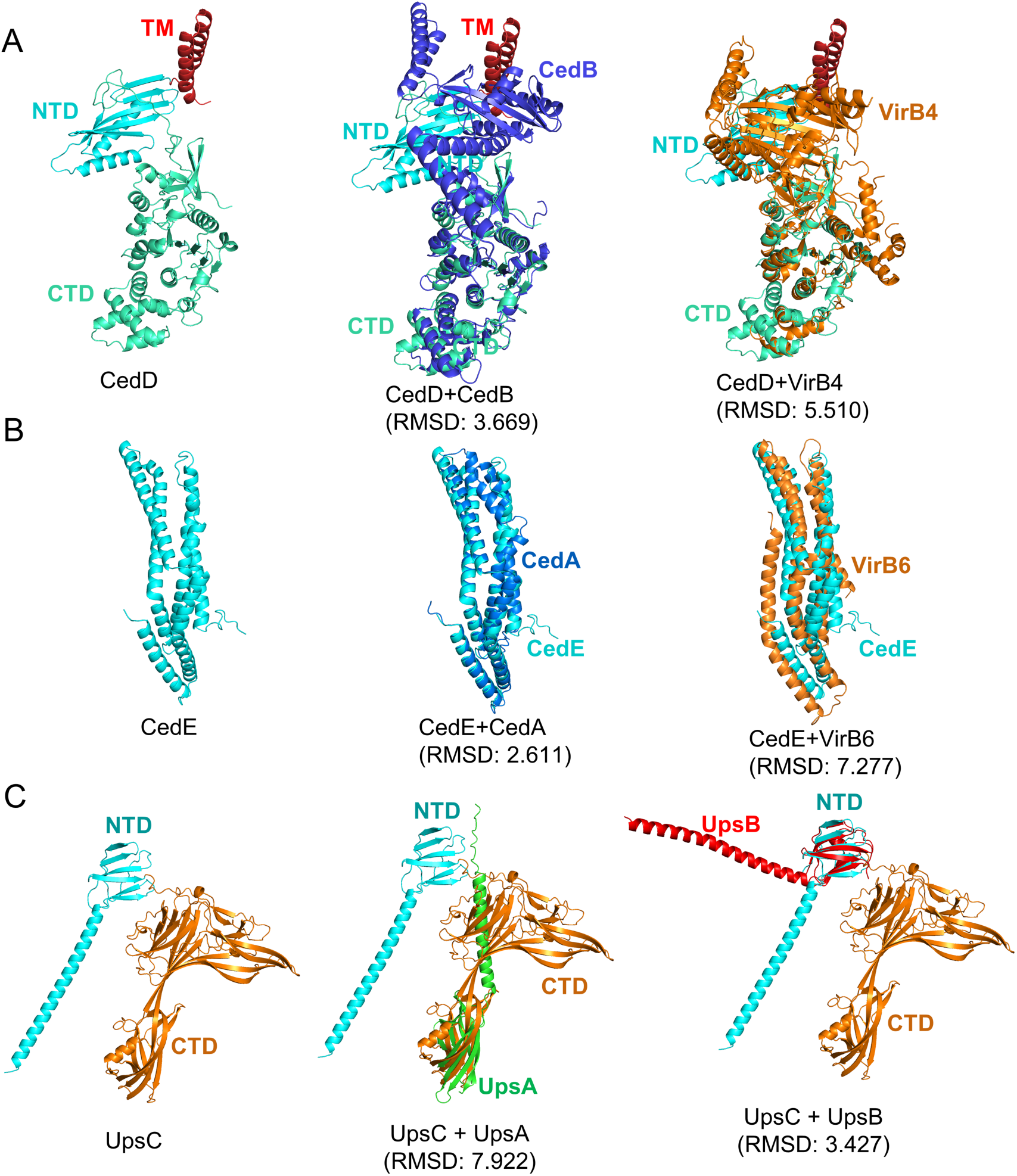
Predicted structures of CedD, CedE, and UpsC and their superposition to the corresponding homologs. (**A**) CedD (Left), CedD to CedB (Middle), CedD to bacterial VirB4 (Right). (**B**) CedE (Left), CedE to CedA (Middle), CedE to bacterial VirB6 (Right). (**C**) UpsC (Left), UpsC to UpsA (Middle), UpsC to UpsB (Right). The structures of CedA, CedB, CedD, CedE, UpsA, UpsB, and UpsC were predicted by AlphaFold server and the structures of VirB4 and VirB6 from the R388 plasmid were generated from 7o41 and 7o3v, respectively. The superpositions were conducted by PyMOL using cealign. The root mean square deviation (RMSD) values are shown.

To confirm *cedD*, *cedE*, and *upsC* were induced by DNA damage, RT-qPCR was conducted using the samples taken at 0 h, 3 h, 6 h, 12 h, and 24 h after 100 J/m^2^ UV treatment (Supplementary Figure 2). At 3 h after UV treatment, the transcriptional levels of *cedD*, *cedE*, and *upsC* were upregulated about 23, 78, and 45 folds, respectively, which were close to those of their corresponding Ced and Ups component genes, *cedB*, *cedA*, and *upsA*. The transcriptional levels of these genes slightly decreased at 6 h and 12 h, and finally returned to the untreated levels at 24 h (Supplementary Figure 2). These results imply that *cedD* and *cedE* function in DNA transport while *upsC* functions in cell contact as their corresponding homologs during DNA damage response.

### Development of a CRISPR-Cas based chromosomal DNA exchange assay

Given that CedA and CedB are involved in DNA import (van Wolferen *et al*., 2016), we speculated that CedD and CedE may play roles in DNA exchange (DNA export and/or import) in cooperation with the known Ced components or in an independent manner. To discriminate whether a gene is involved in DNA export or import, we developed a CRISPR-Cas based chromosomal DNA export and import assay modified from a DNA import assay method reported previously (Liu *et al*., 2020) (Figure 2). Briefly, a target gene and a protospacer sequence within it were selected, and the deletion strain of the target gene was constructed. In the DNA export assay, *lacS* gene was selected as a target and the plasmid pT*lacS* which contains *pyrEF* and a mini-CRISPR array targeting *lacS* was constructed. The pT*lacS* plasmid was transformed into the cells of the receptor strain E233, which contains the complete *lacS* gene. After 1 h incubation to recover, the receptor cells were mixed with the cells of the donor stain E233S, or its derived strains in which the *lacS* genes were absent. If the donor cell can export DNA, the CRISPR-Cas induced double-strand breaks (DSBs) in the receptor cells will be repaired by homologous recombination and the cells will survive on MTAV medium lacking uracil (Figure 2A). The DNA export efficiency is calculated by the CFU of the tested strain in DNA export assay divided by the transformation efficiency of E233 using a non-targeting plasmid. The relative DNA export efficiency is calculated as the DNA export efficiency of the deletion strain divided by that of the control E233S.

**Figure 2.**
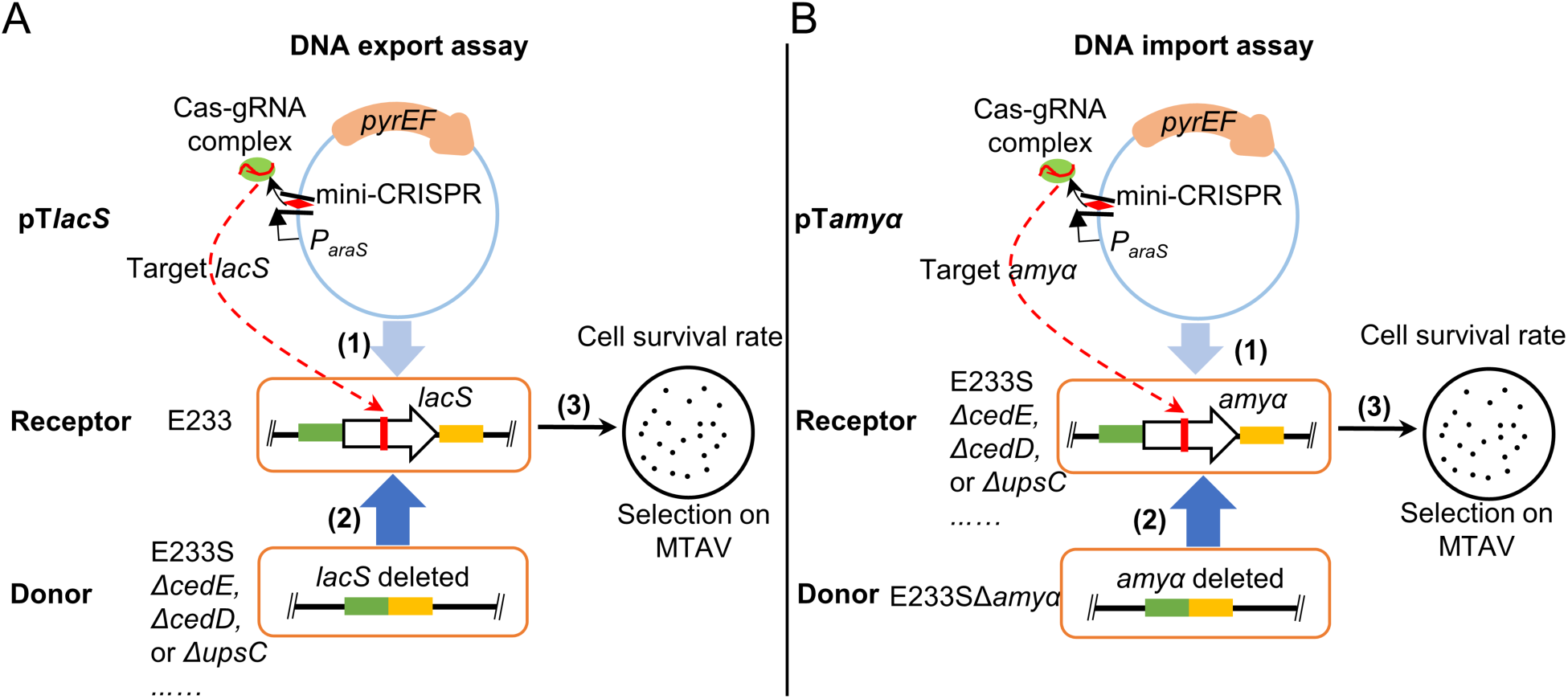
Schematic diagrams showing the established CRISPR-Cas based chromosomal DNA export (A) and import (B) assays. For DNA export assay, the target plasmid pT*lacS* containing a mini-CRISPR array against the *lacS* gene is electro-transformed into E233 containing *lacS* and incubated at 75℃ for 1 h (1). Then, the cells are mixed with each of the donor strains, E233S (*lacS* deleted) or E233S derivatives, Δ*cedD*, Δ*cedE*, or Δ*upsC*, etc. The mixtures are incubated in the inducible medium MTAV at 75℃ for 2 h when the donor chromosomal DNA is transferred into the receptor strain (2). In the cell of the receptor strain, the *lacS* gene is targeted by CRISPR-Cas system resulting in double-strand breaks (DSBs). The receptor cells can be rescued by homologous recombinational repair by the chromosomal DNA (with *lacS* deletion) that is exported from E233S and survive on MTAV plates (3). The donor cell cannot survive due to lack of uracil synthesis ability. Similar strategy is used for the DNA import assay in which *amyα* is used as the target. The DNA export or import efficiency is defined as CFU divided by the transformation efficiency of each strain. Finally, the relative DNA export and import efficiency of different knockout strains is normalized to the efficiency of E233S as 100 %.

For the DNA import assay, the *amyα* gene in E233S was selected as a target and its targeting plasmid pT*amyα* was constructed (Figure 2B). The plasmid was transformed into the receptor strain E233S or its derived strains, which has the complete *amyα*. After incubation, the receptor cells were mixed with the cells of the donor strain, E233SΔ*amyα* (lacking the target gene *amyα*). If a receptor cell can import DNA, the CRISPR-Cas induced DSBs will be repaired, and the cells will survive (Figure 2B). The DNA import efficiency is calculated by the CFU of the tested strain in DNA import assay divided by the transformation efficiency of itself using a non-target plasmid. The relative DNA import efficiency is calculated as the DNA import efficiency of the deletion strain of the Ced and Ups systems divided by that of the control E233S.

To verify if the assay method works, we tested the colony formation of E233 transformed with pT*lacS* (E233::pT*lacS*) and E233S transformed with pT*amyα* (E233S::pT*amyα*) without addition of the donor strains. As shown in Supplementary Figure 3, none, or only one colony formed on these plates, while hundreds colonies formed with the addition of the corresponding donor strains (E233::pT*lacS*×E233S and E233S::pT*amyα*×Δ*amyα*), suggesting that the targeting efficiency of pT*lacS* and pT*amyα* are almost 100 % and most of the colonies were those with repaired DSBs using the untargeted templates from donor strains. In addition, it could be excluded that the donor strains obtained plasmids by natural transformation, since few colonies grew on the plates when E233S cells were directly mixed with the cells of pT*lacS* or E233SΔ*amyα* were directly mixed with pT*amyα* (Supplementary Figure 3). Then, we tested the DNA exchange efficiency of the gene knockout strains of the known Ced and Ups components, individually. Consisted with the previous report, deletion of *cedA* and *cedB* only resulted in decrease of the DNA import efficiency, while deletion of Ups pili operon *upsXEFAB* abolished DNA exchange, possibly by reducing cell aggregation (Figure 3). These results confirmed that our established assay system is efficient for testing DNA export and import.

**Figure 3.**
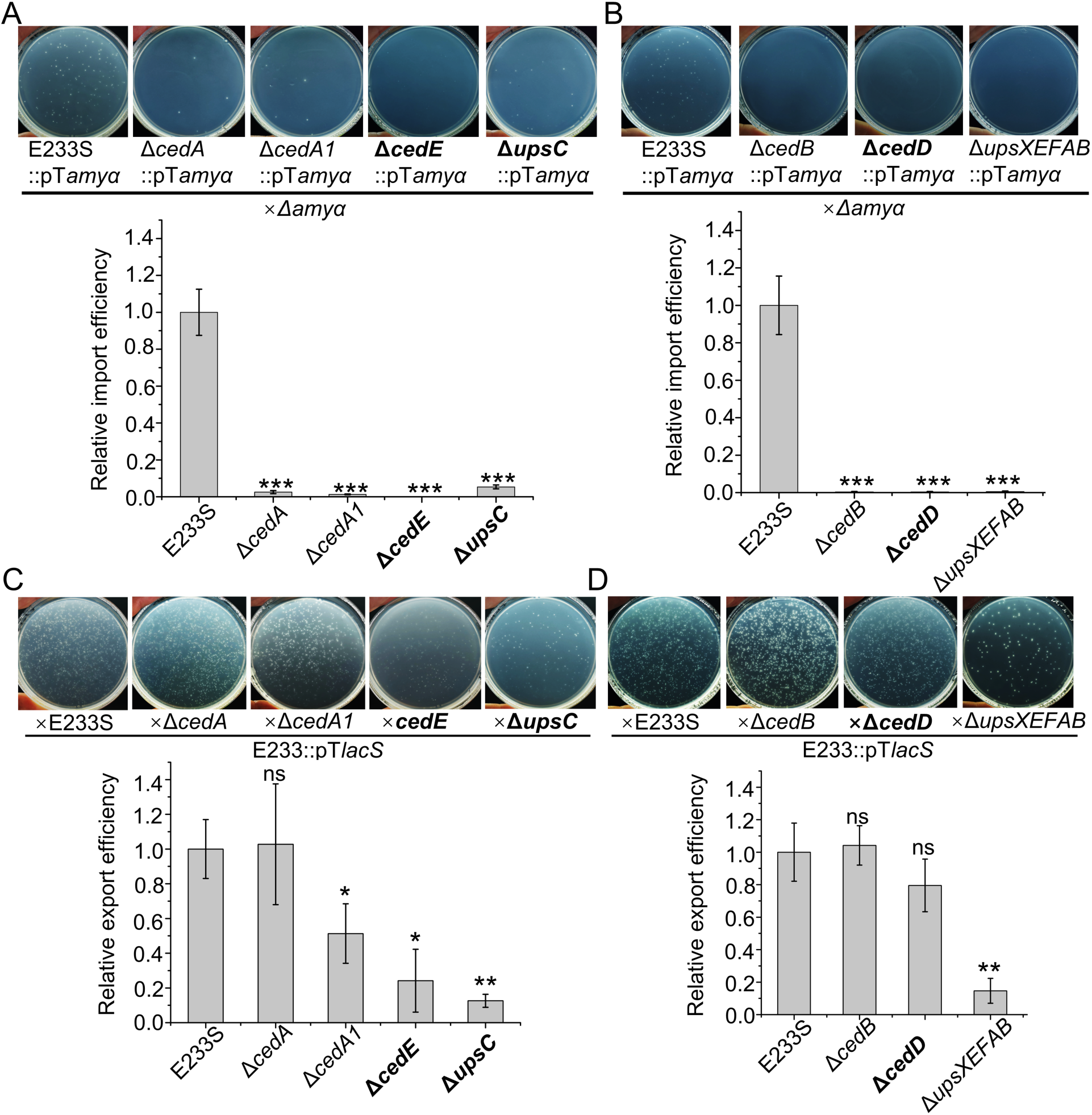
CedE and UpsC are involved in both DNA import and export while CedD is only involved in DNA import. (**A**) Chromosomal DNA import efficiency of Δ*cedA*, Δ*cedA1*, Δ*cedE*, and Δ*upsC*. (**B**) Chromosomal DNA import efficiency of Δ*cedB*, Δ*cedD*, and Δ*upsXEFAB* (deletion of the operon for the Ups pili). (**C**) Chromosomal DNA export efficiency of Δ*cedA*, Δ*cedA1*, Δ*cedE*, and Δ*upsC*. (**D**) Chromosomal DNA export efficiency of Δ*cedB*, Δ*cedD*, and Δ*upsXEFAB*. Representative plates are shown at the upper panels and quantification of the data are shown at the lower panels. The average relative export efficiency was normalized to that of E233S (n=3 independent experiments, ±SD). The significance was calculated by independent samples t-test using SPSS (ns, no significance, *, *P*≤0.05, **, *P*≤0.005, ***, *P*≤0.001).

### CedE and UpsC are critical for both DNA export and import while CedD is only essential for DNA import

Next, deletion strains of genes coding for the new Ced and Ups components were constructed, and their chromosomal DNA exchange ability were examined. We found that the knockout strains Δ*cedD*, Δ*cedE*, and Δ*upsC* lost the DNA import ability, the same as Δ*cedA*, Δ*cedA1*, Δ*cedB*, and Δ*upsXEFAB* (Figure 3A, 3B), suggesting that these three new proteins play essential roles in DNA import, probably synergistically with CedA, CedA1, CedB, and Ups pili. Interestingly, the DNA export efficiency of Δ*cedE* and Δ*upsC* decreased to about 30 % and 10 % of that for E233S, respectively. The DNA export efficiency of Δ*cedA1* also decreased to about 50 %. However, Δ*cedD*, Δ*cedA*, and Δ*cedB* had the similar export efficiency as the control E233S (Figure 3C and 3D). These results suggest that CedE and UpsC together with CedA1 also play important roles in chromosomal DNA export.

### *cedD* and *cedB* encode VirB4-like ATPases and co-occur in crenarchaeal genomes

Previously, CedB and CedD were defined as HerA clade proteins, in which the C-terminal ATPase domain contains synapomorphies including a small residue (typically, glycine) after strand 2, a hydrophobic residue in the α-helix after strand 2, an aspartate in the α-helix immediately after strand 5 (Iyer *et al*., 2004). Here, we found that their N-terminal domains (NTD) are structural homologs of the NTD of VirB4 (Figure 4A), another clade of the FtsK-HerA superfamily functioning in DNA export in T4SS (Atmakuri *et al*., 2004). Interestingly, the NTDs of CedB and CedD are able to superpose to two different parts of the NTD of bacterial VirB4 (Figure 4A) and constitute an intact hypothetical NTD homolog of VirB4. More interestingly, CedB/CedD and VirB4 are distributed in a complementary pattern in Crenarchaea (Figure 4B). CedB and CedD co-occur exclusively in *Acidilobales*, *Sulfolobales*, and most *Desulfurococcales* archaea, while VirB4 were found exclusively in *Fervidicoccales*, *Thermoproteales* and *Desulfurococcus* (Figure 4B), although in *Thermofilales* both CedD and VirB4 (but not CedB) are present. The apparently complementary distribution of CedB-CedD and VirB4 implies that the function of CedB and CedD could be equivalent to that of VirB4, and the T4SS-like DNA exchange system is important for life and therefore conserved in Crenarchaea.

**Figure 4.**
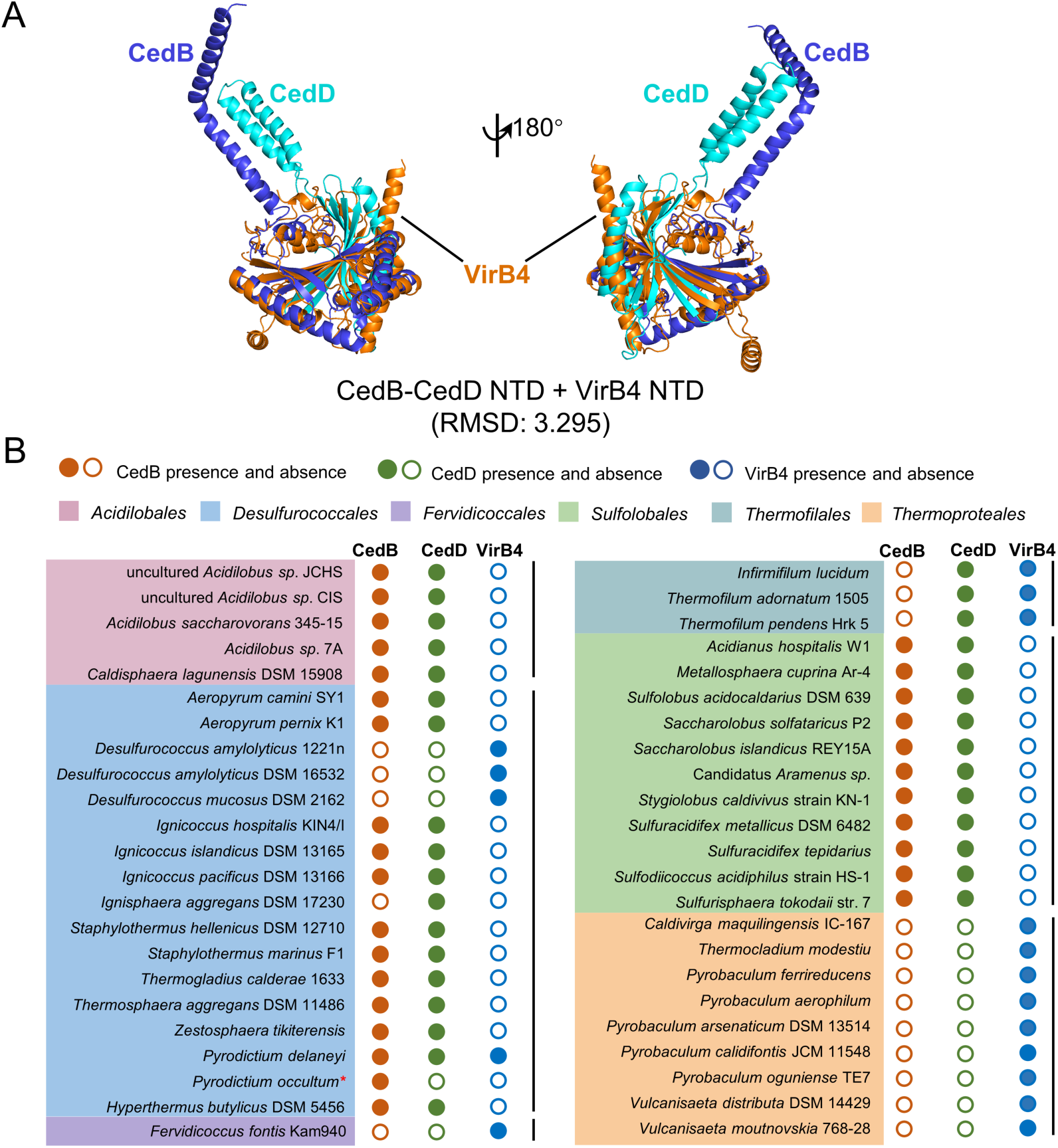
Analysis of the relationship between CedB/CedD and VirB4 in Crenarchaeota. **(A)** The structure of the N-terminal domains (NTDs) complex of CedB-CedD is superposable to VirB4 NTD. The structure of the NTD complex of CedB-CedD was predicted by ColabFold and the structure of the VirB4 NTD from R388 plasmid was generated according to PDB 7o41. The superposition was conducted by PyMOL using Cealign. The RMSD value is shown. **(B)** The distribution of CedB and CedD is complementary to that of VirB4. A total of 85 sequences from 46 species of Crenarchaeota were analyzed. The solid and empty circles indicate presence and absence of the homolog in the analyzed species, respectively. The red asterisk indicates a species of which the complete genome sequence is not available.

Then, we analyzed the phylogenetic relationship of the conserved ATPase domains of CedB, CedD and VirB4. As shown in Supplementary Figure 4, CedB, CedD, and VirB4 homologs were clearly divided into three clades. In the CedB and CedD clades, the homologs from the same orders cluster together, indicating that CedB and CedD possibly evolved from an ancestor ATPase in Crenarchaea and divided at an early evolutionary stage. Supporting this assumption, a VirB4 homolog from *Pyrobaculum calidifontis* clusters with a bacterial VirB4. The homolog could be the oldest VirB4 in Crenarchaeota.

### CedD facilitates the expression of CedB and both form homo-oligomers *in vitro*

To explore the mechanisms of CedB and CedD in DNA import, we attempted to express and purify CedB and CedD proteins in *Sa. islandicus* E233S. Expression of the full length CedB and CedD using strains carrying the pSeSD-based vectors (with D-arabinose inducible promoter P*_araS_*) was unsuccessful. Hence, we tried to express the TMs deleted proteins, CedBΔTM and CedDΔTM using the D-arabinose inducible promoter P*_araS_*, and the full length CedB and CedD using their own promoters induced by DNA damages (Supplementary Figure 5A). Both CedBΔTM and CedDΔTM could be detected when they were co-expressed, while only full length CedD and CedDΔTM were detectable when it was expressed alone (Supplementary Figure 5A). Next, we co-expressed His-tagged CedBΔTM and Flag-tagged CedDΔTM in E233SΔ*cedB*Δ*CedD* using a strain carrying the piDSB-based vector by which the proteins together with DSBs were induced by D-arabinose. The cells were lysed, and the supernatant was subjected to purification with a nickel column and the eluted proteins were collected. The flowthrough of the nickel column was further subjected to purification with anti-Flag magnetic agarose beads. Then the purified samples were analyzed by size exclusive chromatography (SEC). Interestingly, both nickel column purified His-CedBΔTM and anti-Flag magnetic agarose beads purified Flag-CedDΔTM samples exhibited two peaks in the SEC profiles, corresponding to homo-oligomers and monomers in different ratios (Figure 5, Supplementary Figure 5B). Further SEC analysis of the peaks of CedBΔTM showed that the purified CedBΔTM oligomer did not convert to monomer, suggesting that the oligomer was stable, neither did the monomer convert to oligomer, suggesting that monomeric CedBΔTM was stable and formation of its homo-oligomer might need the presence of CedDΔTM (Supplementary Figure 5C and 5D). These results suggest that CedD facilitates the expression and homo-oligomer formation of CedB, although a tight complex of CedB and CedD could not be obtained.

**Figure 5.**
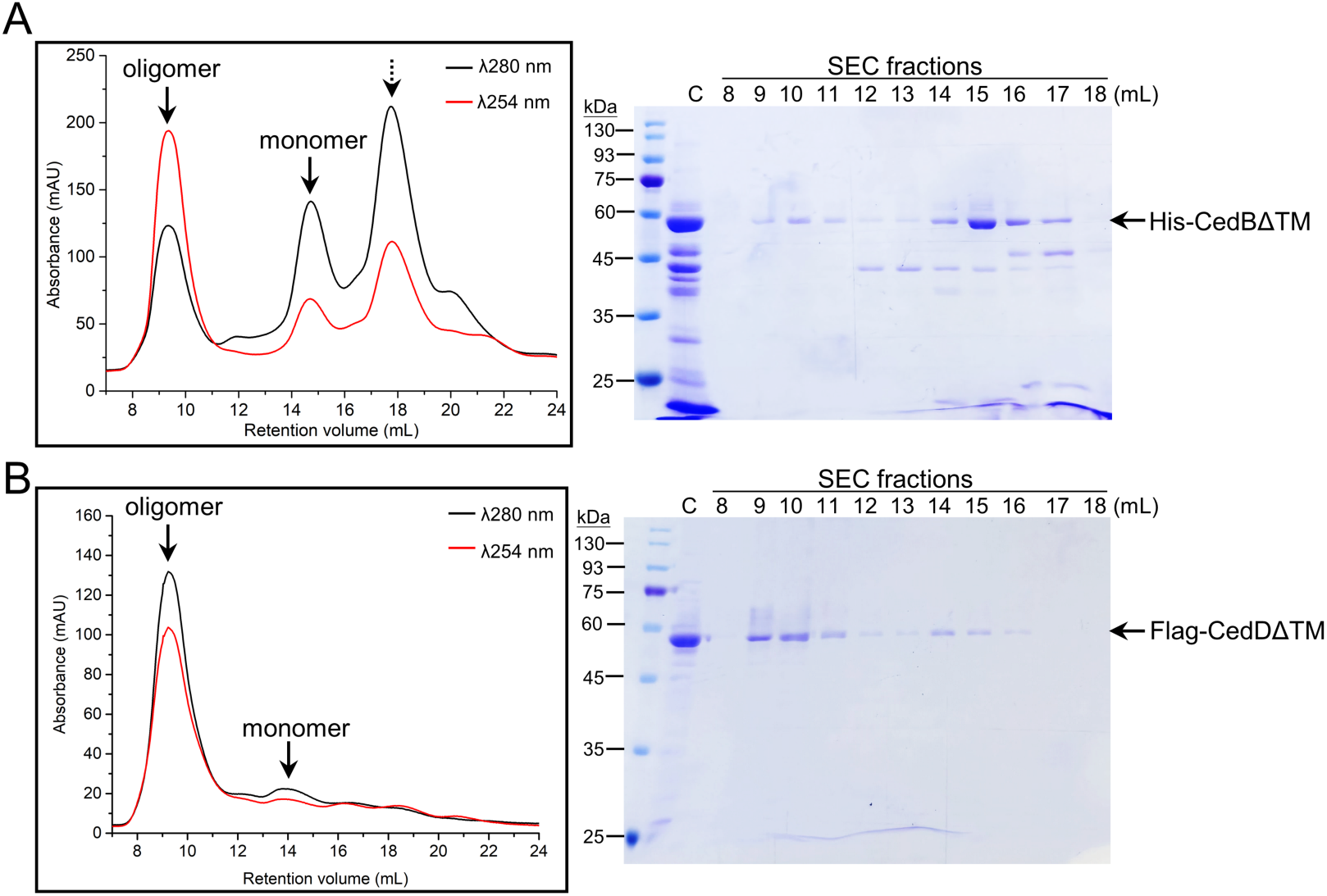
Purified CedB and CedD form homo-oligomers *in vitro*. (**A**) Size exclusive chromatography (SEC) analysis of the nickel column purified His-tagged CedBΔTM (left) and SDS-PAGE analysis of the fractions (right). (**B**) Size exclusive chromatography (SEC) analysis of anti-Flag magnetic agarose purified Flag-tagged CedDΔTM (left) and corresponding SDS-PAGE (right). C, samples before SEC. Lanes 8-18, samples of corresponding fractions (1ml each) in SEC. The theoretical oligomeric and monomeric peaks of CedBΔTM and CedDΔTM are indicated by solid arrows. The peak of the non-specific binding proteins purified with nickel column is indicated by a dotted arrow.

### UpsC functions in cell aggregation in cooperation with Ups

Because UpsC has structural homology to T4P pilin (Figure 1C) and deletion of *upsC* resulted in significant reduction of DNA export and import efficiency (Figure 3), we assumed that UpsC is probably one component of Ups and involved in cell aggregation. To explore the relationship among UpsC and T4Ps, and investigate whether UpsC is involved in cell aggregation in cooperation with the Ups system, we knocked-out genes for the T4Ps in *Sa. islandicus* REY15A, including archaellum (F), adhesion pili (A), putative bindosome (B), and Ups pili (Szabo *et al*., 2007; Zolghadr *et al*., 2007; Frols *et al*., 2008; Henche *et al*., 2012) (Supplementary Figure 6) and constructed a series of deletion strains, including ΔFAB in which the whole archaellum operon, bindosome operon, and *aapE-aapF* operon for adhesion pili assembly were knocked out, Δ*cedA*ΔFAB, Δ*upsC*ΔFAB, and Δ*upsXEFAB*ΔFAB. At 6 h after UV treatment, the strain Δ*cedA*ΔFAB exhibited about 50 % cell aggregation similar to the control ΔFAB, while the cell aggregation ratios of Δ*upsC*ΔFAB and Δ*upsXEFAB*ΔFAB decreased to 20% (Figure 6A). This result supports that UpsC functions in cell aggregation and cooperates with Ups pili while the Ced system is not involved in cell aggregation during DNA damage response (van Wolferen *et al*., 2016). Moreover, UpsC is able to function normally in ΔFAB, suggesting that the assembly of UpsC is independent of archaellum, adhesion pili, and bindosome. We also found that cell aggregation was not fully abolished in the *upsC* or *upsXEFAB* (Figure 6A) and the aggregates in these strains contained only 3 to 7 cells (Figure 6B). This suggests that other factors may play partial roles in cell contact in *Sa. islandicus* REY15A.

**Figure 6.**
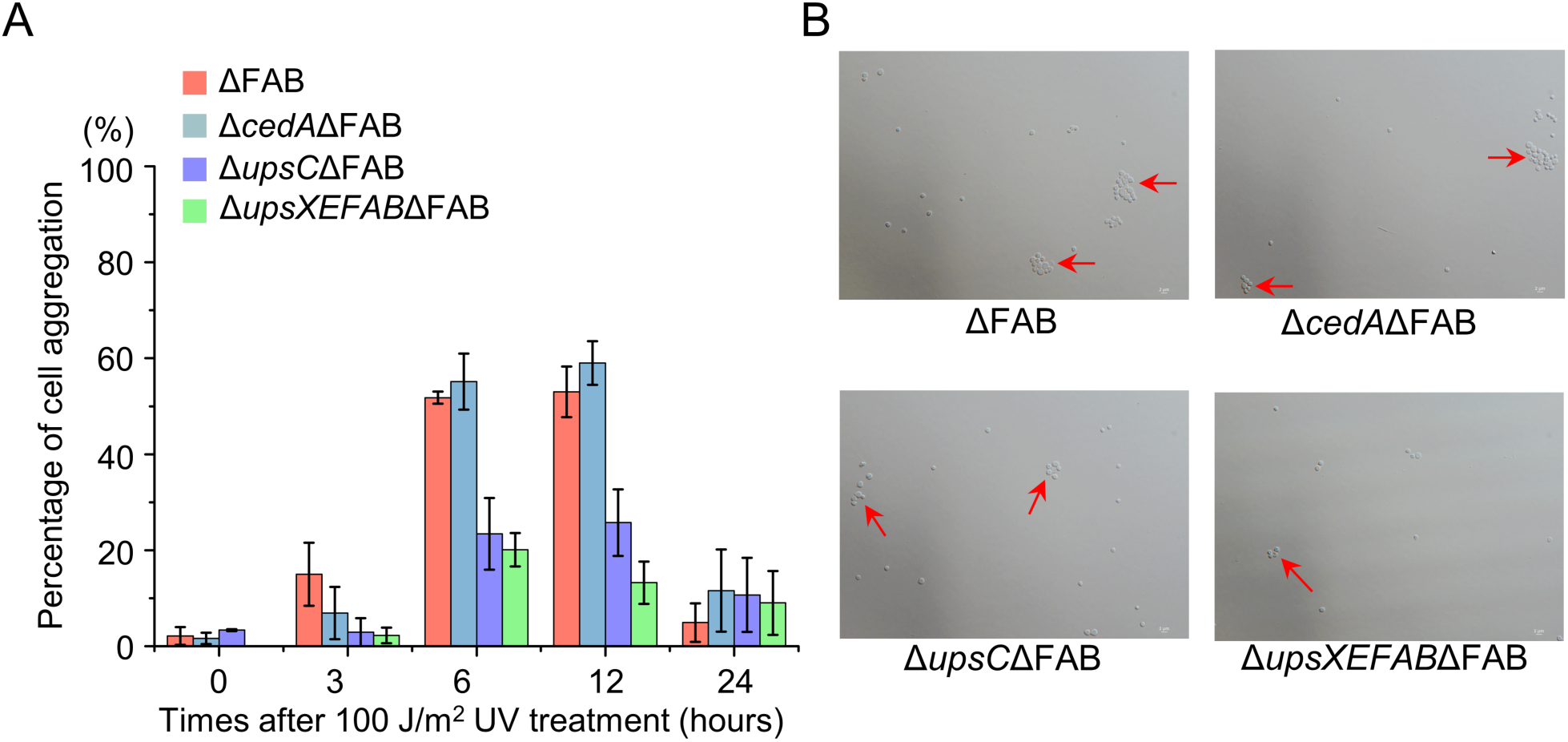
UpsC is involved in UV-induced cell aggregation, which is independent of the archaellum, adhesion pili, bindosome, and Ced systems. (**A**) Percentages of cell aggregation in different strains at 0 h, 3 h, 6 h, 12 h, and 24 h after 100 J/m^2^ UV treatment. The values were based on three independent experiments and the standard deviation (±SD) was indicated. The cell aggregation was defined as cell clustering with at least 3 cells. (**B**) Representative microscopic photos of the cells at 6 h after 100 J/m^2^ UV treatment. The aggregated cells are indicated by red arrows.

## Discussion

DNA export and DNA uptake mechanisms such as conjugation and natural transformation have been studied in detail in bacteria (Johnston *et al*., 2014; Cabezon *et al*., 2015; Waksman, 2019). However, the corresponding mechanisms in archaea are far from clear, perhaps because there are diverse DNA transport pathways and known bacterial homologs such as the relaxase in archaeal conjugative plasmids cannot be identified (Schleper *et al*., 1995; Wagner *et al*., 2017; Catchpole *et al*., 2023). In *Sulfolobales*, the Ced system is involved in DNA import as the DNA damage response mechanism (van Wolferen *et al*., 2016). CedA and CedB are homologs of the T4SS components VirB6 and VirB4, respectively (van Wolferen *et al*., 2016; Beltran *et al*., 2023). The CedA1 pili also shares structural similarity to T4SS pili (Beltran *et al*., 2023). Here, we found that CedD and CedE are additional structural homologs of VirB4 and VirB6, respectively (Figure 1A, 1B, and 4A). These results indicate that the Ced system is a T4SS-like system in Crenarchaeota, suggesting that it could work in a manner similar to bacterial T4SS. Combining the phylogenetic analysis (Supplementary Figures 1 and 4), it seems that the DNA transport system Ted in *Thermoproteales* is the most simplified T4SS while the Ced system in *Sulfolobales* gained more functionally differentiated paralogs during evolution (Beltran *et al*., 2023).

To detect the DNA export and import process individually, we developed a CRISPR-Cas based chromosomal DNA exchange assay (Figure 2). By targeting two protospacers, we could generate DSBs efficiently at single sites by the endogenous CRISPR-Cas systems than the previously reported (Liu *et al*., 2020). In the DNA import assay, almost no colony grow on the plates of Δ*cedB*::pT*amyα*×Δ*amyα*, indicating that the plasmids in the receptor cell did not transfer to the donor. This fact also indicates that the DNA transport from the defined donor to receptor is examined in our method while combination of two directions transfer is examined in the chromosomal marker exchange method (van Wolferen *et al*., 2016). One question remains about how chromosomal and plasmid DNA is discriminated by the cells. On the other hand, the receptor cells, in which the genomic DSBs are induced, are obtained by electroporation, which possibly make the cells in an abnormal status. Hence, we attempted to construct a plasmid which could be used to strictly and efficiently induce DSBs by the CRISPR-Cas system. However, we only obtained the plasmid piDSB which could be strictly regulated but its targeting efficiency is not 100 %. Besides, the measured DNA transport efficiency is not only resulted from the transport processes across the membrane, but also resulted from the DNA processing and homologous recombination processes. In another word, the export and import efficiency is a general reflection of the efficiency in DNA processing, export, import, and homologous recombination. Therefore, to correctly describe the functions of DNA transport proteins like CedB and CedD, homologous recombination efficiency may need to be checked in the deletion strains.

As expected, the components of Ced system are essential for DNA import. Interestingly, two components CedA1 and CedE also function in DNA export (Figure 3). In *Aeropyrum pernix* K1, the CedA1 pili was reported to have the potential to transport DNA (Beltran *et al*., 2023). While in bacteria, the T4SS pili not only serves as a conduit for ssDNA transfer, but also facilitates cell-cell contact by the adhesin VirB5 on the tips of pili (Aly & Baron, 2007; Backert *et al*., 2008; Goldlust *et al*., 2023; Beltran *et al*., 2024). Therefore, the participation of CedA1 and CedE in DNA export suggests that CedA1 of the receptor cell likely forms a DNA conduit which is stabilized by CedA1 and CedE proteins of the donor cell. The fact that both CedA and CedE are essential for DNA import implies that CedA and CedE possibly form a hetero oligomer on cell membrane. CedA2 is a small membrane protein containing two TM helixes like CedA1 and was proposed to form a complex with CedA and CedA1 (van Wolferen *et al*., 2016). In *A. pernix* K1 and *Pyrobaculum calidifontis*, in which the structure of CedA1/TedC pili have been solved, CedA2 is not absent. We tried to construct a *cedA2* deletion strain but failed, possibly the protospacer selected for targeting was not efficient. Therefore, the role of CedA2 needs further investigation.

In bacterial T4SS, apart from the pili and pili assembly components, there are also outer membrane core (OMCC) and inner membrane core complexes (IMCC) (Mace *et al*., 2022). The OMCC was not found in Ced system possibly due to the lack of out membrane in archaea (Albers & Meyer, 2011). In the IMCC, VirB3 and VirB8 connect VirB6 channel on membrane and the ATPase ring of VirB4 in cytoplasm (Hu *et al*., 2019; Mace *et al*., 2022). The homologs of VirB3 and VirB8 are absent in archaea. Instead, about half of the CedB/CedD/VirB4 proteins in the Ced system contain TM helixes, implying that there is direct interaction between CedA/CedE and CedB/CedD. Additionally, the distribution of CedB/CedD is complementary to that of VirB4/TedB in Crenarchaeota, suggesting that CedB and CedD together could function as VirB4 (Figure 4). One hypothesis is that the CedB and CedD assemble into hexamer of dimers like the bacterial VirB4. Homo-oligomers of the TM-deleted CedB and CedD were obtained but purification of a stable hetero-oligomer of TM-deleted CedB and CedD failed (Figure 5). It was reported that overexpressed and purified VirB_3-10_ complex from the pKM101 plasmid contained two side-by-side hexamers of VirB4, quite different from that in the structure *in situ* of the complete pKM101 T4SS in which the VirB4 assembled as hexamer of dimers (Khara *et al*., 2021). These observations suggest that full length CedB and CedD or the whole Ced complex needs to be purified in order to reveal the true structural assemble and mechanism.

In bacteria, the T4SS encoded by the conjugative plasmid is sufficient for DNA transport, while in *Sulfolobales*, the species-specific aggregation mediated by Ups pili is also necessary for DNA exchange in addition to the Ced system (Ajon *et al*., 2011). Here, we identified a new T4P pilin-like protein UpsC. The deletion of other three type T4Ps in *Sa*. *islandicus* REY15A did not inhibit cell aggregation while the deletion of *upsC* or *ups* operon did (Figure 6), suggesting that UpsC is an assembly component of Ups pili. In addition, although *upsC* is not close to the *ups* operon in genome, it co-occur with the *ups* operon in *Sulfolobales* except *Acidianus*. However, the absence of Ups pili-coding genes in other Crenarchaeota implies that cell aggregation is not necessary for DNA transport in these species, or the function of Ups pili is replaced by other cell appendages like archaeal bundling pili found in *Pyrobaculum calidifontis* (Wang *et al*., 2022).

Based on the results of this study and previous reports, we propose an expanded DNA exchange model in *Sa. islandicus* REY15A (Figure 7). When DNA damage occurs, the transcription of DNA damage response genes is induced, such as those coding for the components of the Ups and Ced systems. For cell-cell recognition and aggregation, the Ups pilins, UpsA, UpsB, and UpsC are processed and assembled to form Ups pili to mediate species-specific aggregation (Frols *et al*., 2008; van Wolferen *et al*., 2013; van Wolferen *et al*., 2020). During DNA transport, the chromosomal DNA is firstly processed by nucleases and helicases including the ParB-domain containing nuclease (van Wolferen *et al*., 2015). Then the processed DNA is exported by unknown ATPases from the putative TM channels formed by CedE. Through the conduit of pili formed by CedA1 and possibly CedA2 (Beltran *et al*., 2023), the donor DNA is imported by the oligomers of CedB and CedD ATPase from TM channels formed by CedA and CedE (van Wolferen *et al*., 2016). Finally, the DNA is used for homologous recombination repair in the receptor cells.

**Figure 7.**
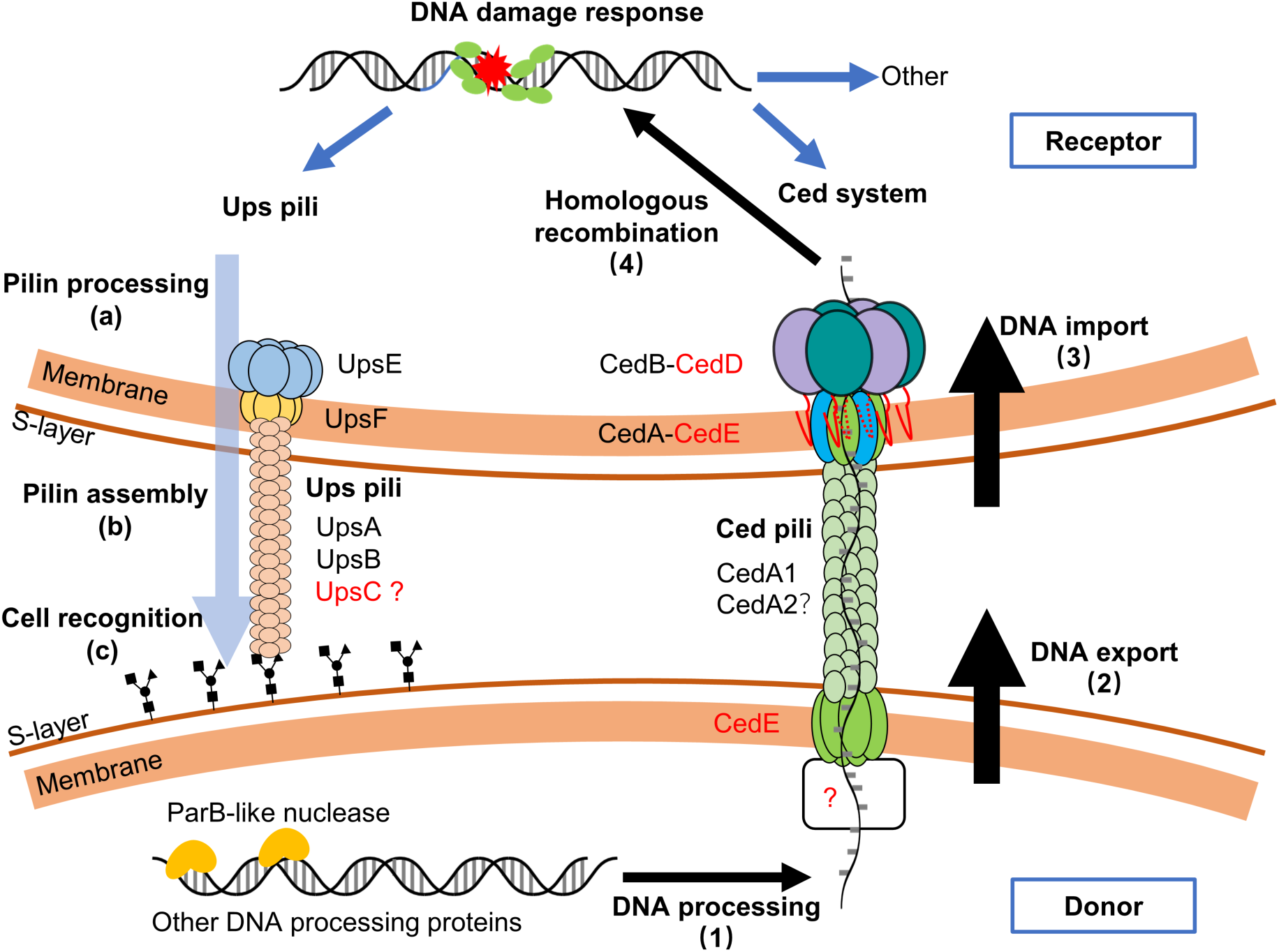
An extended model for DNA exchange in *Sa. islandicus* REY15A. After DNA damages occur in the cell, Ups pilins are processed and assembled to form Ups pili for cell recognition and aggregation. During DNA transfer, the dsDNA is processed by ParB-like nuclease (Saci_1498 homolog) and other DNA processing proteins (1). Then, the processed ssDNA is exported from CedE to CedA1 pili (2). Next, the DNA is imported trough the CedA-CedE transmembrane channel with the assistance of CedB-CedD ATPases (3). Finally, the donor DNA is used for repairing the DNA damages via homologous recombination (4). The new components found in this study are labelled in red.

## Data Availability

All data supporting the findings of this study are available within the article and the Supporting Information, or from the corresponding author upon reasonable request.

## Author Contributions

PW: Investigation, Methodology, Conceptualization, Data analysis, Writing – original draft. MZ: Investigation. YK: Investigation. SL: Methodology, Writing – original draft. JN: Funding acquisition, Writing – review & editing. QH: Supervision, Conceptualization, Funding acquisition, Writing – original draft. YS: Funding acquisition, Supervision, Conceptualization, Writing – review & editing.

## Acknowledgments

We thank members of CRISPR and Archaea Biology Research Center at Shandong University for helpful discussions. We thank technicians from the Core Facilities for Life and Environmental Sciences, State Key Laboratory of Microbial Technology of Shandong University for the assistance in the microscopical experiments. This work was supported by grants from the National Natural Science Foundation of China [32393973] and [32370033] to Y.S., the National Key Research and Development Program of China [2020YFA0906800] to J.N. and Q.H., and the State Key Laboratory of Microbial Technology Open Projects Fund [M2023-20] to Y.S.

