## Supplementary information for "Identification of novel components of the Ced and Ups systems in *Saccharolobus islandicus* REY15A"

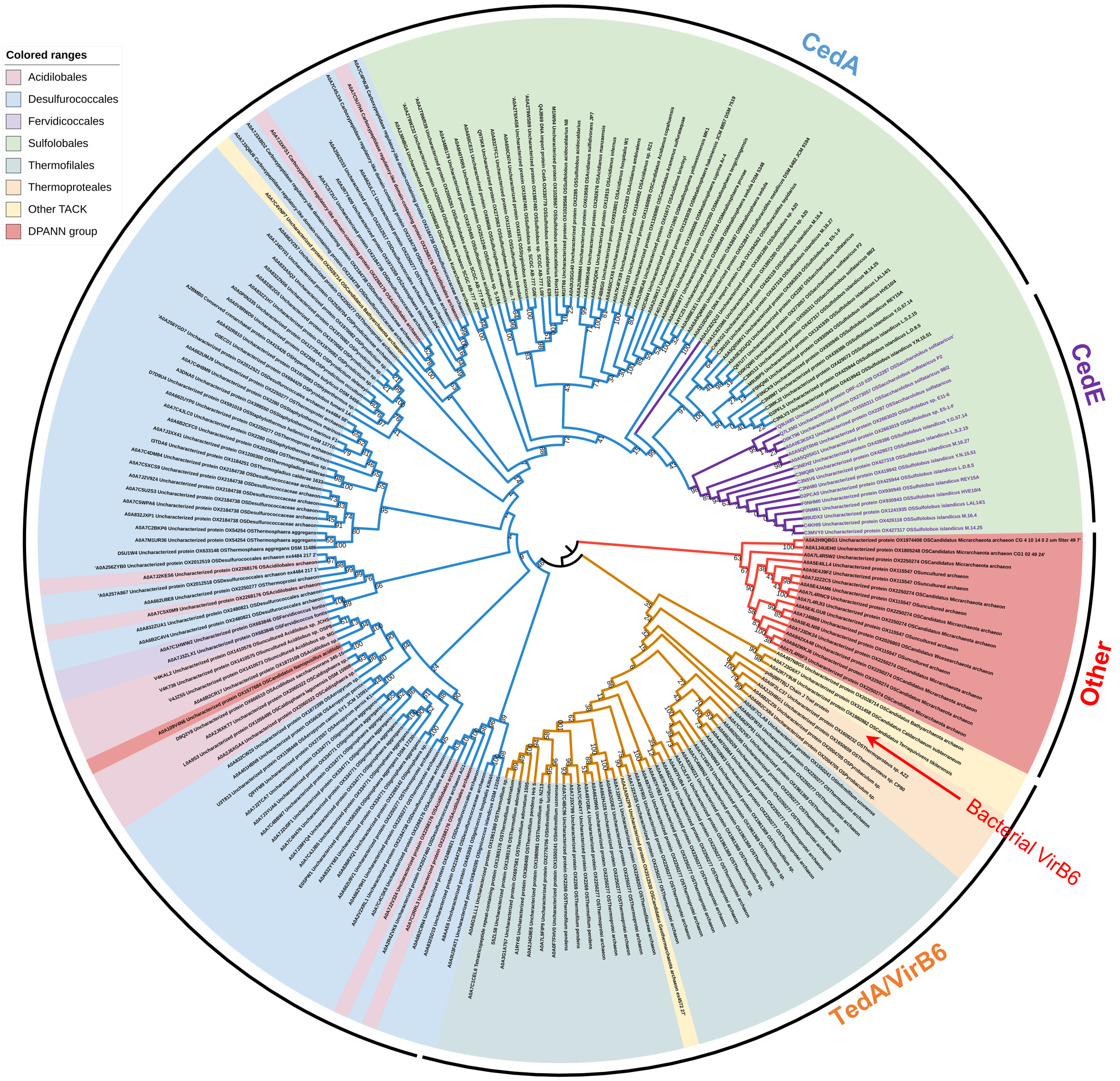


**Supplementary Figure 1. Phylogenetic relationship between archaeal CedA, CedE, TedA and bacterial VirB6.** A total of 209 sequences from archaea and 1 from *Escherichia coli* R388 plasmid were selected, and the amino acid sequence were used to construct the Maximum Likelihood phylogenetic tree by MEGA. The bootstrap value is shown on each branch point. The archaeal protein sequences were downloaded from AFDB clusters, A0A2T9WR39, A0A256ZYB0, and A0A7C2LFF1. The bacterial VirB6 is indicated by red arrow. And the UniProt ID of each protein are shown.

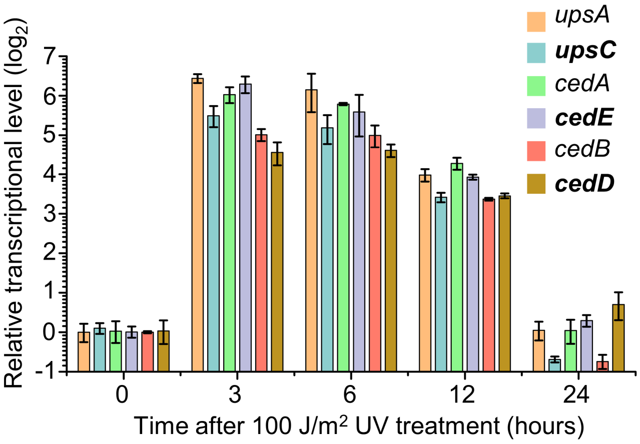


**Supplementary Figure 2. Verification of the transcriptional changes of *cedD*, *cedE*, and *upsC* after UV treatment by RT-qPCR.** The reported Ced and Ups system genes *cedA*, *cedB*, and *upsA* were used as controls. The culture (30 mL) of E233S was treated with 100 J/m$^2$ UV. Samples were taken at 0 h, 3 h, 6 h, 12 h and 24 h after the treatment and subjected to total RNA extraction and RT-qPCR. The transcriptional levels were normalized to the level of *tbp* based on three independent experiments.

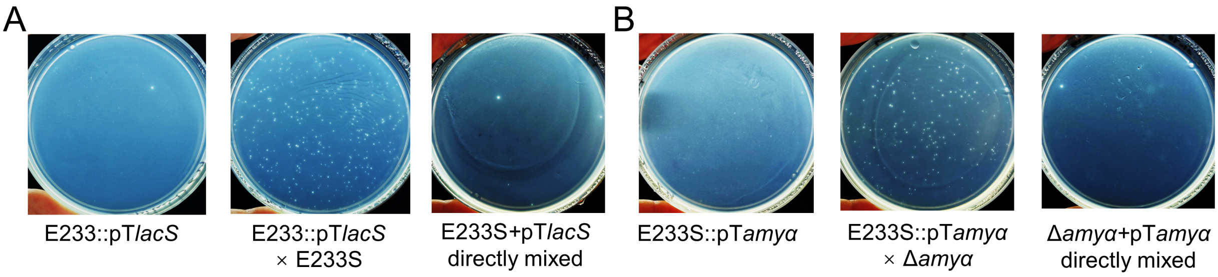


**Supplementary Figure 3. Verification of the CRISPR-Cas based chromosomal DNA export (A) and import (B) assay methods.** Left, plates showing the colony formation of the receptor strains E233 and E233S transformed with the target plasmids pT*lacS* and pT*amyα*, respectively. Middle, the transformed receptor cells mixed and incubated with the donor cells. Right, the donor cells directly mixed with the plasmid. Representative plates are shown.

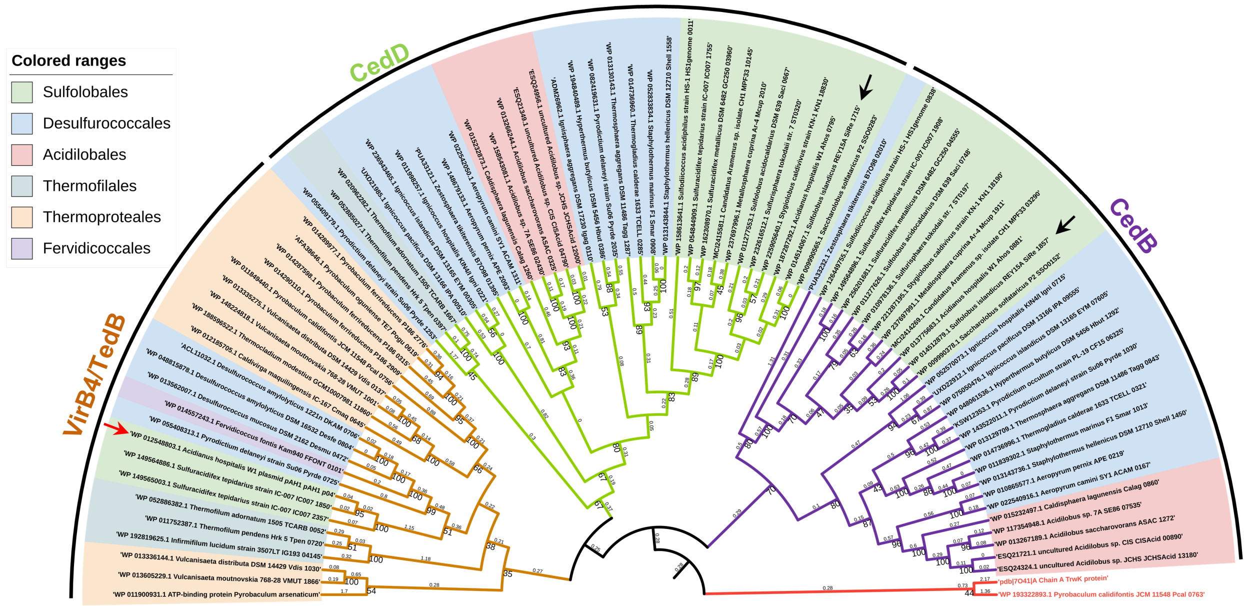


**Supplementary Figure 4. Phylogenetic relationship between CedB, CedD, and VirB4 homologs from Crenarchaeota.** A total of 86 sequences from 47 species were selected and the amino acid sequence of their ATPase domains were used to construct the Maximum Likelihood phylogenetic tree by MEGA. The bootstrap values more than 30 are shown on each branch point and the branch length measured in the number of substitutions per site is indicated on the middle of each branch. The label of VirB4 (PDB: 7o41) from *Escherichia coli* R388 plasmid and *Pyrobaculum calidifontis* are colored in red (bottom right). CedB and CedD from *Sa. islandicus* REY15A are indicated by black arrows and the VirB4 from the conjugative plasmid pAH1 of *Acidianus hospitals* W1 was indicated by the red arrow.

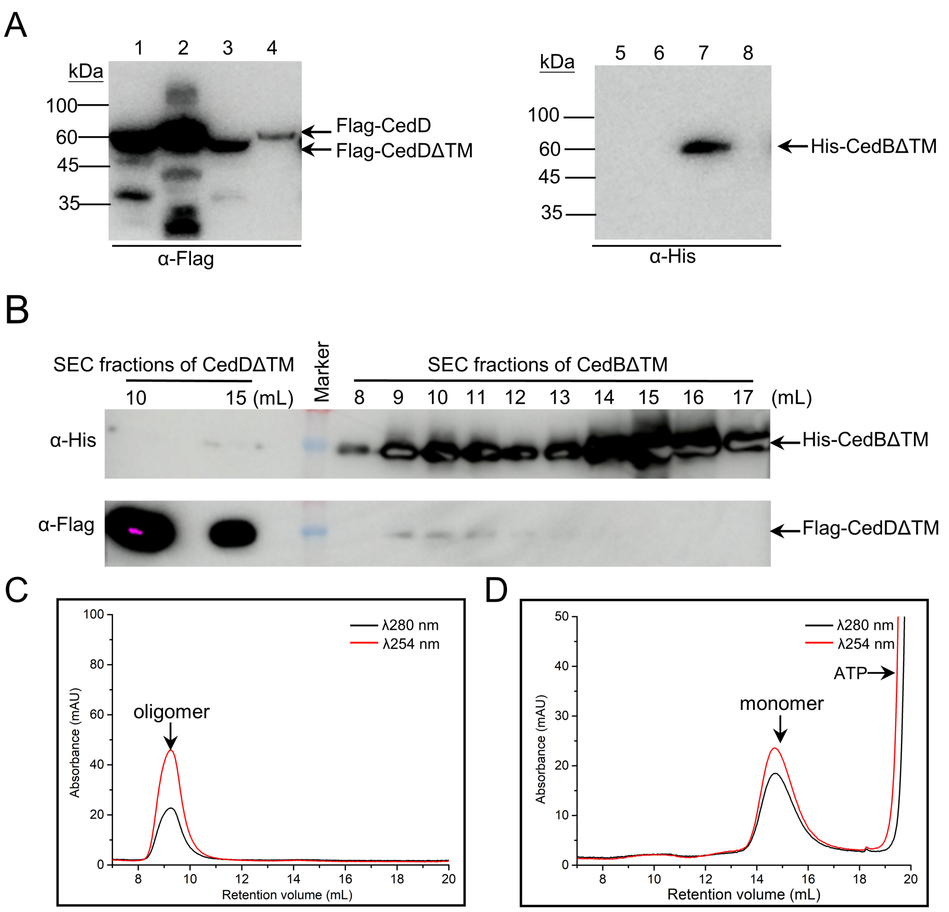


**Supplementary Figure 5. CedD facilitates the expression and homo-oligomer formation of CedB.** (**A**) Western blot of samples from different CedB and CedD expression strains. CedD and CedB were detected with anti-Flag (left) and anti-His antibodies (right), respectively. Lanes 1, E233S/pSeSD-C-Flag-CedDΔTM, D-arabinose; 2, E233S/pSeP$_{cedD}$-C-Flag-CedD, NQO; 3 and 7, E233S/pSeSD-C-His-CedBΔTM/C-Flag-CedDΔTM, D-arabinose; 4 and 8, E233S/pSeP$_{cedB}$-C-His-CedB/P$_{cedD}$-C-Flag-CedD, NQO; 5, E233S/pSeSD-C-His-CedBΔTM, D-arabinose; 6, E233S/pSeP$_{cedB}$-C-His-CedB, NQO. (**B**) Analysis of the size exclusive chromatography (SEC) fractions of CedBΔTM and CedDΔTM in Figure 5 by Western blot using anti-His and and-Flag antibodies. Lanes 8-17, the corresponding SEC fractions of CedBΔTM and CedDΔTM in Figure 5. (**C**) SEC of the purified CedBΔTM oligomer. (**D**) SEC of the purified CedBΔTM monomer. CedBΔTM monomer was incubated with 2.5 mM $MgSO_4$ and 1 mM ATP in 75℃, 20 min before SEC. The arrows indicate the peak of oligomer, monomer, and ATP.

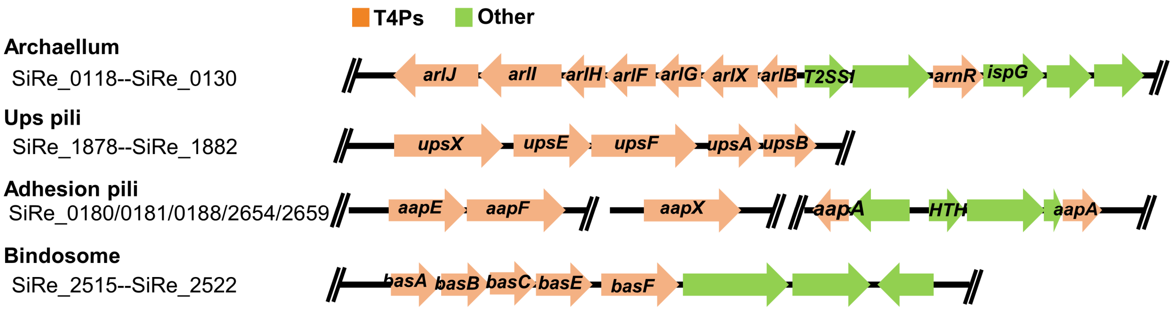


**Supplementary Figure 6. A schematic diagram showing gene context of four type 4 pili systems (T4Ps) in *Sa. islandicus* REY15A.** The pili open reading frames are shown in orange and the others shown in green.

**Supplementary Table 1.** Strains used in this study

| Strain | Properties | Source |
|---|---|---|
| E233 | *Sa. islandicus* REY15A Δ*pyrEF* | (Deng *et al.*, 2009) |
| E233S | *Sa. islandicus* REY15A Δ*pyrEF*Δ*lacS* | (Deng *et al.*, 2009) |
| Δ*amyα* | Deletion of *amyα* in E233S | This study |
| Δ*cedA* | Deletion of *cedA* in E233S | This study |
| Δ*cedA1* | Deletion of *cedA1* in E233S | This study |
| Δ*cedB* | Deletion of *cedB* in E233S | This study |
| Δ*cedD* | Deletion of *cedD* in E233S | This study |
| Δ*cedE* | Deletion of *cedE* in E233S | This study |
| Δ*cedB*Δ*cedD* | Deletion of *cedB* and *cedD* in E233S | This study |
| Δ*upsXEFAB* | Deletion of *upsXEFAB* operon in E233S | This study |
| Δ*upsC* | Deletion of *upsC* in E233S | This study |
| Δ*FAB* | Deletion of the archaellum gene cluster, *aapEaapF* operon, and the bindosome operon in E233S | This study |
| Δ*cedA*ΔFAB | Deletion of *cedA* in Δ*FAB* | This study |
| Δ*upsXEFAB*ΔFAB | Deletion of *upsXEFAB* operon in Δ*FAB* | This study |
| Δ*upsC*ΔFAB | Deletion of *upsC* in Δ*FAB* | This study |

**Supplementary Table 2.** Plasmids used in this study

| Plasmids | Properties | Source |
|---|---|---|
| pGE | *Sulfolobus-E. coli* shuttle vector containing mini-CRISPR and *pyrEF* for CRISPR-Cas based gene editing | (Li *et al.*, 2016) |
| pSeSD | *Sulfolobus-E. coli* shuttle vector containing *araS*-SD promoter and MCS for proteins expressed in E233S | (Peng *et al.*, 2012) |
| piDSB | Insertion of a D-arabinose inducible protospacer at mini-CRISPR array of pGE to generate DNA double stranded break | This study |
| pGE *amyα* KO | For deletion of *amyα* | This study |
| pGE *cedA* KO | For deletion of *cedA* | This study |
| pGE *cedA1* KO | For deletion of *cedA1* | This study |
| pGE *cedB* KO | For deletion of *cedB* | This study |
| pGE *cedD* KO | For deletion of *cedD* | This study |
| pGE *cedE* KO | For deletion of *cedE* | This study |
| pGE *upsXEFAB* KO | For deletion of *upsXEFAB* operon | This study |
| pGE *arl* KO | For deletion of archaellum gene cluster | This study |
| pGE *aapEaapF* KO | For deletion of *aapEaapF* operon | This study |
| pGE *bas* KO | For deletion of bindosome operon | This study |
| pSeSD-C-His-CedBΔTM | To express C-terminal His tagged CedBΔTM by pSeSD | This study |
| pSeSD-C-Flag-CedDΔTM | To express C-terminal Flag-tagged CedDΔTM by pSeSD | This study |
| pSeP$_{cedB}$-C-His-CedB | To express C-terminal His-tagged CedB by replacing P$_{araS}$ promoter on pSeSD to P$_{cedB}$ promoter | This study |
| pSeP$_{cedD}$-C-Flag-CedD | To express C-terminal Flag-tagged CedD by replacing P$_{araS}$ promoter on pSeSD to P$_{cedD}$ promoter | This study |
| pSeSD-C-His-CedBΔTM /C-Flag-CedDΔTM | To co-express C-terminal His-tagged CedBΔTM and C-terminal Flag-tagged CedDΔTM by pSeSD | This study |
| pSeP$_{cedB}$-C-His-CedB /P$_{cedD}$-C-Flag-CedD | To co-express C-terminal His tagged CedB and C-terminal Flag-tagged CedD by replacing P$_{araS}$ promoter on pSeSD to P$_{cedB}$ and P$_{cedD}$ promoters, respectively | This study |
| piDSB-P$_{araS}$-C-His-CedBΔTM /P$_{araS}$-C-Flag-CedDΔTM | To co-express C-terminal His tagged CedBΔTM and C-terminal Flag-tagged CedDΔTM using P$_{araS}$ promoter on piDSB | This study |

**Supplementary Table 3.** Sequences of main oligonucleotides used in this study.

| Oligonucleotides | Sequence (5’ to 3’) |
|---|---|
| *cedA* qPCR-F | CAGATCCACTACTTCAGTTTTCATCTTTT |
| *cedA* qPCR-R | CCTAAAAGGAAGTGATAATAAAAGAATACC |
| *cedE* qPCR-F | CTCTCTTACACCATAGGTACACTACTT |
| *cedE* qPCR-R | GAGGAGGCCAAAGATGTTAGCTAAT |
| *cedB* qPCR-F | GATGGAATATTGTTAGGAAAGGATCC |
| *cedB* qPCR-R | CACATTTAGCAGCGTGGATAGCC |
| *cedD* qPCR-F | TAACGGTGTCAGAGATAACCTACAC |
| *cedD* qPCR-R | TGTTATGTCCTTTACTCTTATACCTACC |
| *upsA* qPCR-F | GTCAATCCTAATACCGGACAAGCAT |
| *upsA* qPCR-R | GCCAGGTTGTAAGAGGTCGTTATTA |
| *upsC* qPCR-F | ATGCTAATCAAGTCTTTGTGGGAGTC |
| *upsC* qPCR-R | TACTTAAATGTTGCGTTGTAGACTGGTA |
| *tbp* qPCR-F | CAACAGTTACGTTAGAGCAAAGTTTGG |
| *tbp* qPCR-R | GTAACCTTGGGCTGTTCTAATCTGA |
| pT*lacS* spacer | TTTCAACAGCCTCCATATCTTTCTCCGTCAACGGTTGGAA |
| pT*amyα* spacer | GTATGCTCTTTGTGATGTTCTCCAAAAGTCTCGTAATCTA |
| piDSB spacer | TTGAGGCTAGTTCTTTGAATATTTTTGCTCTAGGATCATA |
| $P_{cedB}$ SphI-F | TGCTGCATGCATTGAAATCCCGGGTAGAACAAAGTATT |
| $P_{cedB}$ MluI-F | ATCGACGCGTATTGAAATCCCGGGTAGAACA |
| CedBΔTM NdeI-F | TATACATATGTATAATATCGAAGATGGAGT |
| CedB noStop SalI-R | ATAAGCGTCGACGAGCGTGCTACTGGCTAA |
| $P_{cedD}$ SphI-F | TGCTGCATGCAGAGTTCAACTATGTTTTTCTTACTTCT |
| CedDΔTM NdeI-F | TATACATATGCATATAAATAAATATATTCT |
| CedD Flag MluI-R | GGAGACGCGTTTATTTATCGTCATCATCTTTATAGTCAGACATTTTTTTAATT TTAATT |

The restriction enzyme sites are underlined.

**Supplementary Table 4. A**nnotation of uncharacterized genes unregulated more than 30 folds after DNA damage reagent (NQO) treatment in *Sa. islandicus* REY15A.

| Gene ID | Gene annotation | Putative functions | mRNA abundance ratio (FPKM) (Sun *et al.*, 2018) | |
|---|---|---|---|---|
| | | | E233S -NQO | E233S +NQO |
| SiRe_0014 | 5'-3' exonuclease NurA-like | DNA processing | 1 | 35.557 |
| SiRe_0020 | Flg_new_2 domain-containing protein | Cell contact | 1 | 91.566 |
| SiRe_0137 | Major facilitator transporter | MFS transporter | 1 | 129.008 |
| SiRe_0187 | Coiled-coil containing protein | DNA binding | 1 | 124.001 |
| SiRe_0269 | Hypothetical protein | Unknown | 1 | 46.608 |
| SiRe_0426 | ABC transporter related membrane transporter | ABC transporter | 1 | 61.611 |
| SiRe_0589 | DUF1156 domain containing, putative adenine-specific DNA methylase | DNA processing | 1 | 108.693 |
| SiRe_0670 | SWIM (SWI2/SNF2 and MuDR) Zinc finger domain containing protein | Unknown | 1 | 54.801 |
| SiRe_0936 | Glycosyl hydrolase 15 | Unknown | 1 | 41.046 |
| SiRe_1040 | Hypothetical protein | Unknown | 1 | 38.389 |
| **SiRe_1715** | **VirB4 like ATPase** | **DNA transport** | **1** | **141.412** |
| **SiRe_1957** | **Prepilin domain-containing protein** | **Cell contact** | **1** | **128.999** |
| **SiRe_2100** | **CedA paralog** | **DNA transport** | **1** | **73.161** |
| SiRe_2101 | Acyl-CoA dehydrogenase | Lipid metabolism | 1 | 31.776 |
